# Modularity and Morphometrics: Error Rates in Hypothesis Testing

**DOI:** 10.1101/030874

**Authors:** Guilherme Garcia, Felipe Bandoni de Oliveira, Gabriel Marroig

## Abstract

The study of modularity in morphological systems has increased in the past twenty years, parallel to the popularization of geometric morphometrics, which has led to the emergence of different criteria for detecting modularity on landmark data. However, compared to usual covariance matrix estimators, Procrustes estimators have properties that hinder their use. Here, we compare different representations of form, focusing on detecting modularity patterns defined *a priori;* we also compare two metrics: one derived from traditional morphometrics (MHI) and another that emerged in the context of landmark data (RV). Using Anthropoid skulls, we compare these metrics over three representations of form: interlandmark distances, Procrustes residuals, and local shape variables. Over Procrustes residuals, both tests fail to detect modularity patterns, while in remaining representations they show the distinction between early and late development in skull ontogeny. To estimate type I and II error rates, we built covariance matrices of known structure; these tests indicate that, considering both effect and sample sizes, tests using MHI are more robust than those using RV. However, both metrics have low power when used on Procrustes residuals. Thus, we conclude that the influence of development and function is poorly represented on Procrustes estimators for covariance matrices.

## Introduction

Modularity is a characteristic property that biological systems exhibit regarding the distribution of interactions between their composing elements; that is, in a given system, subsets of elements, denominated modules, interact more among themselves than with other such subsets (Newman, 2006; Mitteroecker & Bookstein, 2007; Wagner *et al*., 2007). This property has been well documented at different levels of biological organization, from the dynamics of metabolic networks (e.g. Ravasz *et al*., 2002; Andrade *et al*., 2011) to the structure of interactions among individuals in populations (e.g. Fortuna *et al*., 2008) and among species in ecological communities (e.g. Genini *et al*., 2010).

Regarding morphological systems, the concept of modularity is associated with the framework of morphological integration (Olson & Miller, 1958; Cheverud, 1996), which refers to the organization of covariances or correlations between morphological elements and the hypotheses concerning their relationships. In this context, modularity refers to the uneven distribution of genetic effects over phenotypic variation articulated through development (genotype/phenotype map; Wagner, 1996); in a classical quantitative genetics view, these genetic effects are the result of pleiotropy and linkage disequilibrium (Falconer & Mackay, 1996; Lynch & Walsh, 1998). A genotype/phenotype map composed of clusters of genes that affect clusters of traits (with little overlap) exhibits a modular organization; this structure is thought to emerge as the result of selection for distinct demands (Wagner & Altenberg, 1996; Espinosa-Soto & Wagner, 2010; Rueffler *et al*., 2012; Melo & Marroig, 2015). For instance, the decoupling between fore- and hindlimb function in certain mammalian lineages such as bats (Young & Hallgrímsson, 2005) and apes (Young *et al*., 2010) is associated with the modularization of both structures, as shown by reduced phenotypic correlations between fore- and hindlimbs and increased correlations between elements within these limbs.

The recognition of variational modules (Wagner & Altenberg, 1996; Wagner *et al*., 2007) using covariance or correlation patterns in adult populations involves an uderstanding of the underlying developmental and functional dynamics among morphological traits (Polly, 2008; Zelditch & Swiderski, 2011). Skull development in mammals is composed of a series of steps, such as neurocranial growth induced by brain development, and growth mediated by muscle-bone interactions, with spatiotemporal overlapping between such steps (Hallgrímsson & Lieberman, 2008; Herring, 2011; Cardini & Polly, 2013). Both timing and scope of each step is regulated by different profiles of genetic expression exhibited by cells originated from different embryonic precursors and their response to signaling factors expressed at the regional level; the response to these signaling factors further changes cell expression profiles, thus generating a feedback loop of diferentiation (Turing, 1952; Marcucio *et al*., 2005; Meinhardt, 2008; Hallgrímsson *et al*., 2009; Franz-Odendaal, 2011; Minelli, 2011). Each step in this temporal hierarchy may be regarded as modular, since they affect a coherent subset of tissues more so than others (Hallgrímsson & Lieberman, 2008), although each step affect adjoining regions through interactions among developing tissues (Cheverud *et al*., 1992; Lieberman, 2011; Esteve-Altava & Rasskin-Gutman, 2014); thus, the overlapping of such processes throughout development may complicate their association with correlation patterns (Hallgrímsson *et al*., 2009).

Furthermore, variation in growth rates, which emerges into size variation (Pélabon *et al*., 2013; Porto *et al*., 2013), has a particular importance in the context of mammalian morphological systems. Here size variation refers to variation in both scale (isometric variation) and scale relationships (allometric variation). This source of variation affects the overall level of correlations between morphological traits (Wagner, 1984; Young & Hallgrímsson, 2005), and the magnitude of integration has important consequences for both the evolution of mean phenotypes (Schluter, 1996; Marroig & Cheverud, 2005, 2010; Cardini & Polly, 2013) and the evolution of morphological integration itself (Oliveira *et al*., 2009; Porto *et al*., 2009, 2013; Shirai & Marroig, 2010).

In this context, adaptive landscapes may be the central component governing both the stability and divergence in integration patterns, as both stabilizing (Jones, 2007; Arnold *et al*., 2008) and directional (Jones *et al*., 2012; Melo & Marroig, 2015) selection have been shown to produce changes in integration patterns. The empirical evidence available demonstrates that both stability (e.g. Marroig & Cheverud, 2001; Oliveira *et al*., 2009; Porto *et al*., 2009; Willmore *et al*., 2009) and divergence (e.g. Monteiro & Nogueira, 2010; Grabowski *et al*., 2011; Sanger *et al*., 2012; Haber, 2015) of integration patterns are possible outcomes of the evolutionary process. Therefore, the question of whether any of these two scenarios is the rule or exception at macroevolutionary scales remains open, although some theoretical and methodological differences between these works with respect to the representation of morphological features need to be taken into consideration.

### Morphometries

Traditionally, morphological features are measured using distances among elements defined in general terms, such as “cranial length” or “cranial width”. Pearson & Davin (1924) introduced the notion that measurements should be restricted to single osteological elements, preferably as distances between homologous features that could be identified in a wide taxonomic coverage. Cheverud (1982) accomodates this notion into the framework of morphological integration, thus considering individual measurements over single bones as local representations of regional phenomena, that is, the functional, developmental and genetic interactions that produce covariances among these elements. The influence of such interactions over covariance or correlation matrices can be accessed by defining a subset of measurements which fall under the scope of a particular process and estimating a metric that summarizes such partitioning; the null hypothesis that this partition is undistinguishable from randomly-defined partitions can then be tested using Monte Carlo methods (Mantel, 1967; Cheverud *et al*., 1989).

In the past three decades, geometric morphometrics (Bookstein, 1982, 1991; Kendall, 1984; Rohlf & Slice, 1990; Goodall, 1991) have been consolidated as a quantitative framework for the representation of biological shape as geometric configurations of homologous features (landmarks; Bookstein, 1991). Two principles are central here: first, the conceptual and statistical separation of size and shape as components of biological form; second, the use of superimposition-based methods (GPA: Generalized Procrustes Analysis; Rohlf & Slice, 1990; Goodall, 1991) for the estimation of shape statistical parameters, such as mean shape and shape covariance structure. Procrustes estimators were proposed as a solution to the problem that landmark configurations are arbitrarily rotated and translated; such nuisance parameters are impossible to estimate without any assumptions (Goodall, 1991; Lele & McCulloch, 2002), a situation known as the identifiability problem (Neyman & Scott, 1948).

Although the use of Procrustes estimators is currently widespread in geometric morphometrics toolboxes (e.g. Klingenberg, 2011; Adams & Otárola-Castillo, 2013), it has not been without criticisms, either with respect to the estimation of mean shape configurations (e.g. Lele, 1993; Kent & Mardia, 1997; Huckemann, 2012) and of shape covariance matrices (Walker, 2000; Adams *et al*., 2004; Linde & Houle, 2009; Márquez *et al*., 2012). For configurations in two-dimensional space, Procrustes estimators for mean shape perform well under isotropic landmark covariance structure (Kent & Mardia, 1997), a situation of null covariances among landmarks and coordinates; however, when this assumption does not hold, Procrustes estimates for mean shape behave badly, especially when shape variation is high (Huckemann, 2011). The example provided by Linde & Houle (2009) demonstrates that when such assumption is broken shape covariance patterns are also poorly estimated; if the unknown landmark covariance matrix is structured due to regional differences in covariance-generating processes, such variation will be displaced and effectively spread out through the entire landmark configuration.

A number of alternatives for estimating shape covariance matrices have already emerged; some of these alternatives (e.g. Monteiro *et al*., 2005; Theobald & Wuttke, 2006; Linde & Houle, 2009; Zelditch *et al*., 2009) propose modifications to the Procrustes analysis in order to deal with heterogeneity in landmark covariance structure. Márquez et al. (2012) propose another definition of shape descriptors using interpolation-based techniques (Cheverud & Richtsmeier, 1986; Bookstein, 1989) as a starting point; such descriptors refer to infinitesimal expansions or retractions in reference to a unknown mean shape (Woods, 2003) estimated at definite locations amidst sampled landmarks. The authors argue that these descriptors are proper local measurements of shape variation, as they can directly be linked to biological processes that generate covariation among morphological elements.

Despite these caveats regarding Procrustes estimators for covariance matrices, the use of such estimators for investigating aspects of morphological integration has increased in the past ten years (e.g. Klingenberg *et al*., 2004; Drake & Klingenberg, 2010; Goswami & Polly, 2010; Martínez-Abadías *et al*., 2011; Sanger *et al*., 2012). The results found by these authors are sometimes in stark constrast with similar works using interlandmark distances (e.g. Cheverud *et al*., 1997; Oliveira *et al*., 2009; Porto *et al*., 2009). For instance, Martínez-Abadías et al. (2011) found a pattern of strong integration among partitions in human skull covariance patterns, while Oliveira et al. (2009) and Porto et al. (2009) demonstrated that humans are among the most modular examples of mammalian skull covariance patterns. Likewise, while Cheverud et al. (1997) had shown that 70% of the pleiotropic effects are confined to either anterior or posterior mandibular components, Klingenberg et al. (2004) has found no evidence for a modular distribution of pleiotropic effects among the partitions of the mouse mandible using the same strain of intercrossed mice at the same generation. Furthermore, aside from the issues regarding Procrustes estimators, these works also propose different methods to quantify the effects of interactions over covariance patterns. For example, Martínez-Abadías et al. (2011) and Sanger et al. (2012) use the RV coefficient, a multivariate correlation coefficient defined by Escoufier (1973) which has been used to quantify modular relationships over landmark covariance patterns since Klingenberg (2009) has proposed its use in this context.

### Objectives

In the present work, we compare the methods described by Cheverud et al. (1989) and Klingenberg (2009) to test *a priori* defined modularity patterns using anthropoid primates as a model organism. In order to compare the performance of these methods with respect to different representations of form, individuals in our sample are represented both as interlandmark distances and shape variables. Furthermore, we used an approach based on the construction of theoretical covariance matrices; such matrices are used in order to estimate Type I and Type II error rates for both methods. Since these methods were designed under different frameworks, the present work puts some effort into unifying both methods into the same conceptual and statistical framework, in order to produce meaningful comparisons.

## Methods

### Sample

The database we used here (Table 1) consists of 21 species, distributed across all taxonomic ranks within Anthropoidea above the genus level. We selected these species from a broader database (Marroig & Cheverud, 2001; Oliveira *et al*., 2009) in order to reduce the effects of low sample sizes over estimates of modularity patterns. Individuals in our sample are represented by 36 registered landmarks, measured using a Polhemus 3Draw (for Platyrrhini) and a Microscribe 3DS (for Catarrhini). Twenty-two unique landmarks represent each individual (Figure S1, Table S1), since 14 of the 36 registered landmarks are bilaterally symmetrical. For more details on landmark registration, see Marroig & Cheverud (2001) and Oliveira et al. (2009).

**Table 1:**
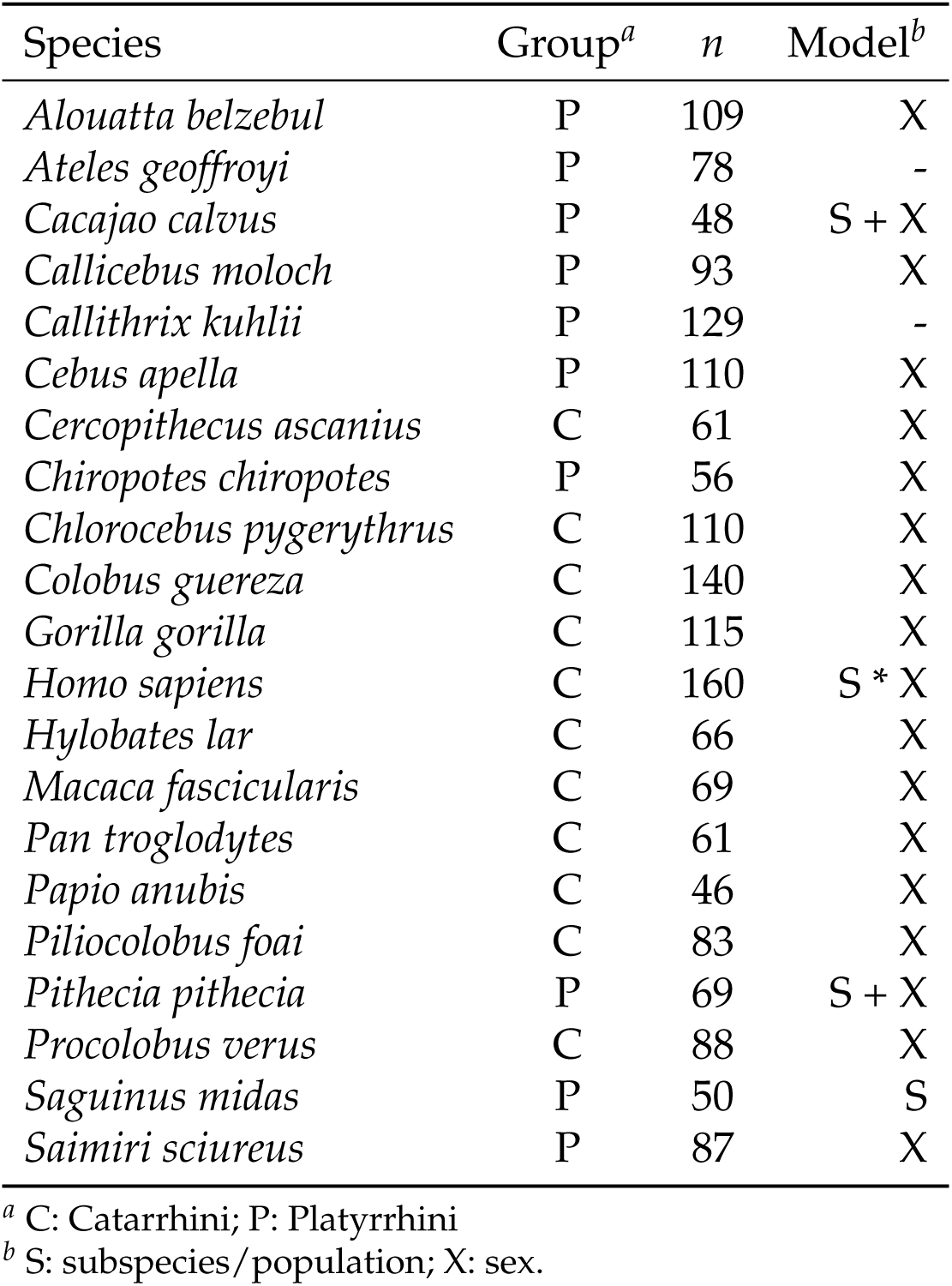
Twenty-one species used in the present work, along with sample sizes and linear models adjusted.

For each OTU, we estimated phenotypic covariance and correlation matrices for three different types of variables: tangent space residuals, estimated from a Procrustes superimposition for the entire sample, using the set of landmarks described on both Table S1 and Figure S1 (henceforth Procrustes residuals); interlandmark distances, described in Table S2; and local shape variables (Márquez *et al*., 2012), which are measurements of infinitesimal log volume transformations between each sample unit and a reference (mean) shape, based upon an interpolation function that describes shape variation between sampled landmarks. In this context, we used thin plate splines as interpolating functions (Bookstein, 1989). We obtained 38 transformations corresponding to the locations of the mipoints between pairs of landmarks used to define interlandmark distances, in order to produce a dataset that represents shape (i.e., form without isometric variation; Bookstein, 1991; Zelditch *et al*., 2004) while retaining the overall properties of the interlandmark distance dataset, such as dimensionality for example. Furthermore, we were able to use the same hypotheses of trait associations for both types of variables since the position of local shape variables through the skull mirrors the position of interlandmark distances, although they are conceptually different types of measurements.

Here we considered only covariance or correlation structure for the symmetrical component of variation; therefore, prior to any analysis, we controlled the effects of variation in assymmetry. For interlandmark distances, we averaged bilateral measurements within each individual. For both Procrustes residuals and local shape variables, we followed the procedure outlined in Klingenberg et al. (2002) for bilateral structures by obtaning for each individual a symmetrical landmark configuration, averaging each actual shape with its reflection along the sagittal plane; we estimate local shape variables afterwards. With respect to Procrustes residuals, landmarks placed along the sagittal plane will have zero variation in the direction normal to this plane; we aligned all specimens’ sagittal plane to the *xz* plane, thus removing the *y* component for each of these landmarks from covariance/correlation matrices.

For each dataset, we estimated covariance and correlation matrices after removing fixed effects of little interest in the present context, such as sexual dimorphism, for example. For interlandmark distances and local shape variables these effects were removed through a multivariate linear model adjusted for each species, according to Table 1; for Procrustes residuals, the same effects were removed by centering all group means to each species’ mean shape since the loss of degrees of freedom imposed by the GPA prohibits the use of a full multivariate linear model over this kind of data to remove fixed effects.

In order to consider the effects of size variation on modularity patterns we used different procedures to remove the influence of size from each type of variable. For interlandmark distances we used the approach established by Bookstein et al. (1985); if **C** is a correlation matrix, we obtained a correlation matrix **R** without the effect of size using the equation

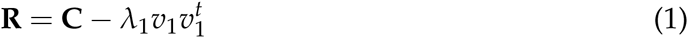

where *λ*_1_ and *υ*_1_ refer respectively to the first eigenvalue and eigenvector of the spectral decomposition of **C**, since this eigenvector commonly represents size variation in mammals, especially when interlandmark distances are considered (Wagner, 1984; Mitteroecker *et al*., 2004; Mitteroecker & Bookstein, 2007); *t* denotes matrix transpose.

For Procrustes residuals and local shape variables the effects of isometric variation were removed by normalizing each individual to unit centroid size. However, allometric relationships still influence covariance or correlation structure. In order to remove this effect we used a procedure based upon Mitteroecker et al. (2004), which relies on the estimation of an allometric component *a* for each OTU, composed of normalized regression coefficents for each of the *m* shape variables (either Procrustes residuals or local shape variables) over log Centroid Size. If **S** is a covariance matrix, we obtained a covariance matrix **R** without the influence of allometric relationships using the equation

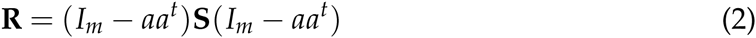

where *I_m_* represents the identity matrix of size *m*. Therefore, our empirical dataset consists of six sets of covariance/correlation matrices, corresponding to each type of morphometric variables considering the presence or absence of size variation.

### Empirical Tests

Using these six sets of covariance/correlation matrices, we tested the hypotheses of trait associations described in Table S1 for Procrustes residuals and Table S2 for interlandmark distances and local shape variables. These trait sets are grouped with respect to their scope; two regional sets (Face and Neurocranium), each divided into three localized trait sets (Oral, Nasal and Zygomatic for the Face; Orbit, Base and Vault for the Neurocranium).

For all hypotheses, we estimated Modularity Hypothesis Indexes (MHI; Porto *et al*., 2013) and the RV coefficient (Klingenberg, 2009). Both statistics are estimated by partitioning covariance or correlation matrices into blocks; if **A** is a covariance or correlation matrix, the partition

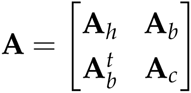

indicates that the block **A***_h_* contains covariances or correlations between traits that belong to the trait set being considered, while **A***_c_* represents the complementary trait set; **A***_b_* represents the block of covariances or correlations between the two sets. Thus, covariance (**S**) or correlation (**C**) matrices can be partitioned into a similar scheme. We estimated MHI values using the equation

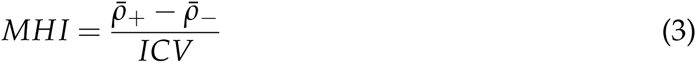

where 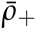 represents the mean correlation in *Ch*, 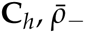 represents the mean correlation in the remaining sets (both **C***_b_* and **C***_c_*), and *ICV* is the coefficient of variation of eigenvalues of the associated covariance matrix, which is a measurement of the overall integration between all traits (Shirai & Marroig, 2010). We estimated RV coefficients for each hypothesis using the relationship

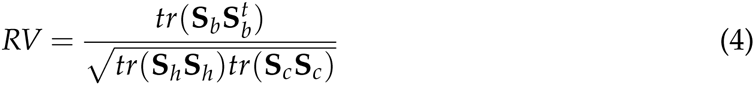

where *tr* represents the sum of diagonal elements in any given matrix (*tr*A = Σ*_i_ a_ij_*). The partitioning scheme outlined above assumes that the complementary trait set does not represent an actual hypothesis; however, we may choose to consider that both sets (**A***_h_* and **A***_c_*) represent two distinct hypothesis. The estimation of RV coefficients remains the same; however, MHI values are estimated considering that 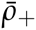 is the average correlation in both **C***_h_* and **C***_c_*, while 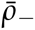 represents the average correlation only in **C***_b_*. In the case of the distinction between Facial and Neurocranial traits, we estimated MHI values in this manner, reporting values for this estimate under the denomination ‘Neuroface’, following Marroig & Cheverud (2001), along independent MHI estimates for each region. Furthermore, since both Face and Neurocranium are two disjoint trait sets when any morphometric variable type is considered, RV coefficient values for either set are equal; therefore, a single RV value is reported for both regions, for each variable type.

In order to test the hypothesis that a trait set represents a variational module, we used a randomization procedure generating 1000 random trait sets with the same number of traits as the original set, calculating MHI and RV values for each iteration. For each trait set and covariance/correlation matrix, we used these values to construct distributions for both statistics representing the null hypothesis that a given trait set is a random arrangement without meaningful relationships; we then compare this null distribution to the real value obtained. For MHIs we consider this null hypothesis rejected when the real value is higher than the upper bound for the distribution, considering the significance level established; for RV coefficients, the null hypothesis is rejected when the real RV value is lower than the lower bound for the distribution, also considering significance level. For Procrustes residuals the randomization procedure maintains coordinates within the same landmark together in each randomly generated trait set, following Klingenberg & Leamy (2001).

While the procedure for estimating significance for MHIs is derived from Mantel’s (1967) approach (as outlined by Cheverud *et al*., 1989), we chose to generate null distributions for MHI directly, instead of estimating matrix correlation values for both real and randomized matrices. Estimated *p*-values in both cases remain the same, and the additional step of calculating matrix correlations would produce an unnecessary difference between the estimation of signficance for MHI and RV.

### Estimation of Error Rates

We used a set of theoretical covariance matrices to investigate Type I and II error rates for either MHI and RV metrics; the construction of such matrices is detailed in the Supplemental Information. Here, it suffices to say that we build two different sets of covariance matrices: one with known modular patterns embedded, referred to as **C***_s_* matrices, and another that represents random covariance structure, denominated **C***_r_* matrices. For each of the six sets of empirical matrices we use here, we built a set of 10000 covariance matrices of each case (either **C***_s_* or **C***_r_*) that mimic the statistical properties of each set, obtaining from these matrices samples of increasing sample size (20,40, 60, 80,100 individuals).

If samples were generated from a **C***_r_* matrix, this represents a situation of a true null hypothesis for either tests, since the correlation matrix used to produce the sample was generated by a permutation of the hypothesis being tested. Therefore, testing hypotheses over **C***_r_* matrices allows us to estimate Type I error rates, or the proportion of cases in which a true null hypothesis is rejected, given a significance level. In an adequate test, we expect that both quantities, significance level and Type I error rate, will be identical.

The opposite case, when we sampled **C***_s_* matrices, represents a situation in which we know that the null hypothesis of either test is false, since we are testing the hypothesis that the partitioning scheme used to generate that particular matrix actually represents two variational modules. Thus, we estimated Type II error rates, or the probability that a false null hypothesis is not rejected, given a significance level; here, we represent Type II error using the power for each test, by simply calculating the complementar probability to Type II error rate. In an adequate test, we expect that power will rapidly reach a plateau when significance level is still close to zero, and further increasing *P*(*α*) will not produce a great increase in power.

Our estimates of power for both statistics should also be controlled for effect size, since sampled correlations may generate a correlation structure that is not detected due to small differences among within-set and between-set correlations. For each correlation matrix sampled, we estimate squared between-set correlations (*b*^2^), in order to use it as an estimate of effect size that is not directly associated with either MHI and RV metrics. We expect that power for either tests decreases with increasing *b^2^* values, as effect size would also decrease.

#### Software

All analysis were performed under R 3.2.2 (R Core Team, 2015). Source code for all analyses can be found at http://github.com/wgar84. Previous tests we made indicated no differences between our estimation of empirical RV coefficients, based upon our own code, and estimates provided by MorphoJ (Klingenberg, 2011). In order to obtain symmetrical landmarks configurations, we used code provided by Annat Haber, available at http://life.bio.sunysb.edu/morph/soft-R.html.

## Results

### Empirical Tests

Tests performed using MHI for localized trait sets (Oral, Nasal, Zygomatic, Orbit, Base, and Vault; Figure 1a) detect a consistent pattern among OTUs for interlandmark distances and local shape variables; in the first set the Oral subregion is detected as a modular partition, and, when size is removed, the Vault subregion is also detected; both Orbit and Base region are not detected in any of these tests. With local shape variables, Oral, Vault Nasal, and Zygomatic sets were detected consistently across OTUs; the removal of allometric variation affects only the detection of the Vault in some groups. Furthermore, the Base sub-region is detected only in 3 of 42 tests performed over local shape variables, pooling together size and size-free correlation patterns.

**Figure 1:**
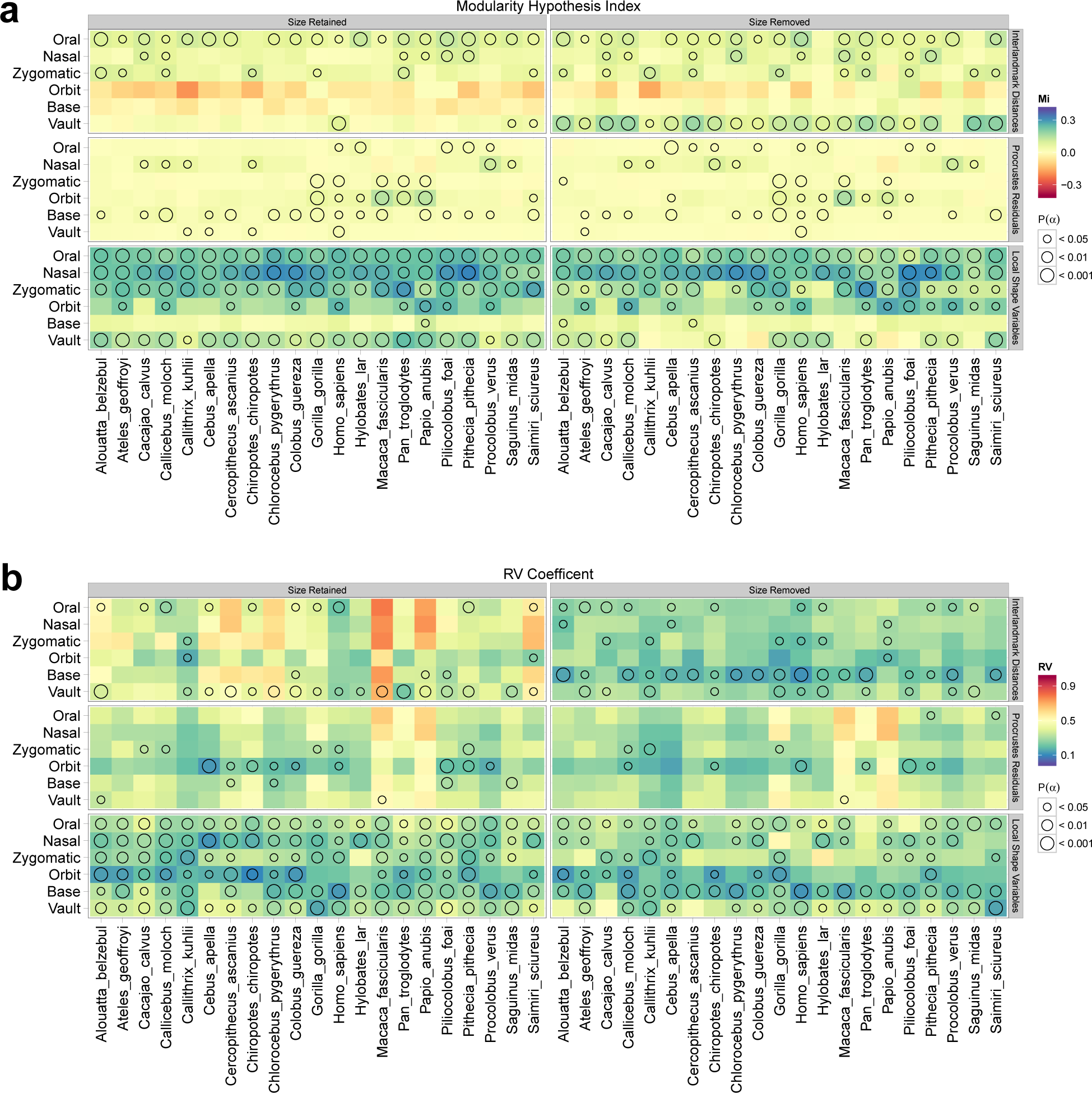
MHI (a) and RV (b) values for localized trait sets. Circles indicate whether a trait set is recognized as a variational module in a given OTU, with *P*(*α*) indicated by the legend. Notice that blue values are associated with higher MHI values and lower RV values, as the alternate hypothesis for each statistic is formulated in a corresponding manner; see text for details.

For Procrustes residuals, the pattern of detection among sub-regions and OTUs is more inconsistent; for instance, the Base sub-region is detected in several OTUs, which contrasts this type of morphometric variable with the other two types. Observing the actual MHI values, Figure 1a also indicates that Procrustes residuals display a low variance of this metric within each OTU, while interlandmark distances and local shape variables display a consistent pattern of variation, with lower values for the Base and, for interlandmark distances, Orbit trait sets, while the Oral, Nasal and Vault regions display higher values consistently.

Tests performed using RV coefficients (Figure 1b) show a more irregular pattern for each variable type. When interlandmark distances are considered, most tests detect the Vault sub-region with size variation retained, and the Base sub-region when size variation is removed. For Procrustes residuals, few tests are able to reject their null hypotheses, detecting only a handful of valid modular partitions. Tests performed on local shape variables display the opposite behavior: almost all partitions are detected, regardless of whether allometric variation has been retained or removed. Moreover, RV values display a pattern of marked variation among OTUs, more so than between values within each OTU; notably, *Macaca fascicularis* and *Papio anubis* show RV values much higher than those estimated on remaining species. Such pattern can be observed both on interlandmark distances with size retained and in Procrustes residuals.

With respect to regional trait sets (Face and Neurocranium), tests performed using MHI (Figure 2a) indicate a pattern consistent with the findings regarding localized sets (Figure 1a). Considering interlandmark distances, Facial traits are detected as a valid modular partition both with size variation retained or removed, while Neurocranial traits are detected as a valid partition only when size variation is removed. This pattern mirrors the contrast between Oral and Vault traits in the localized sets regarding interlandmark distances. For Procrustes residuals, Neurocranial traits are a valid partition with both size retained and removed; once again, this pattern mirrors the detection of the Basicranial partition as valid in the localized sets. Finally, in local shape variables, both Face and Neurocranium are detected as valid with size retained; with size removed, only the Face is recognized consistently. The same can be observed for localized sets, where removing allometric variation affects the detection of the Vault set in some OTUs. The tests for the distinction of within-set and between-set correlations for these two sets (designated ‘Neuroface’) show a pattern that is consistent with tests for the individual sets: if one of the sets was previously detected, this distinction is also detected as valid.

**Figure 2:**
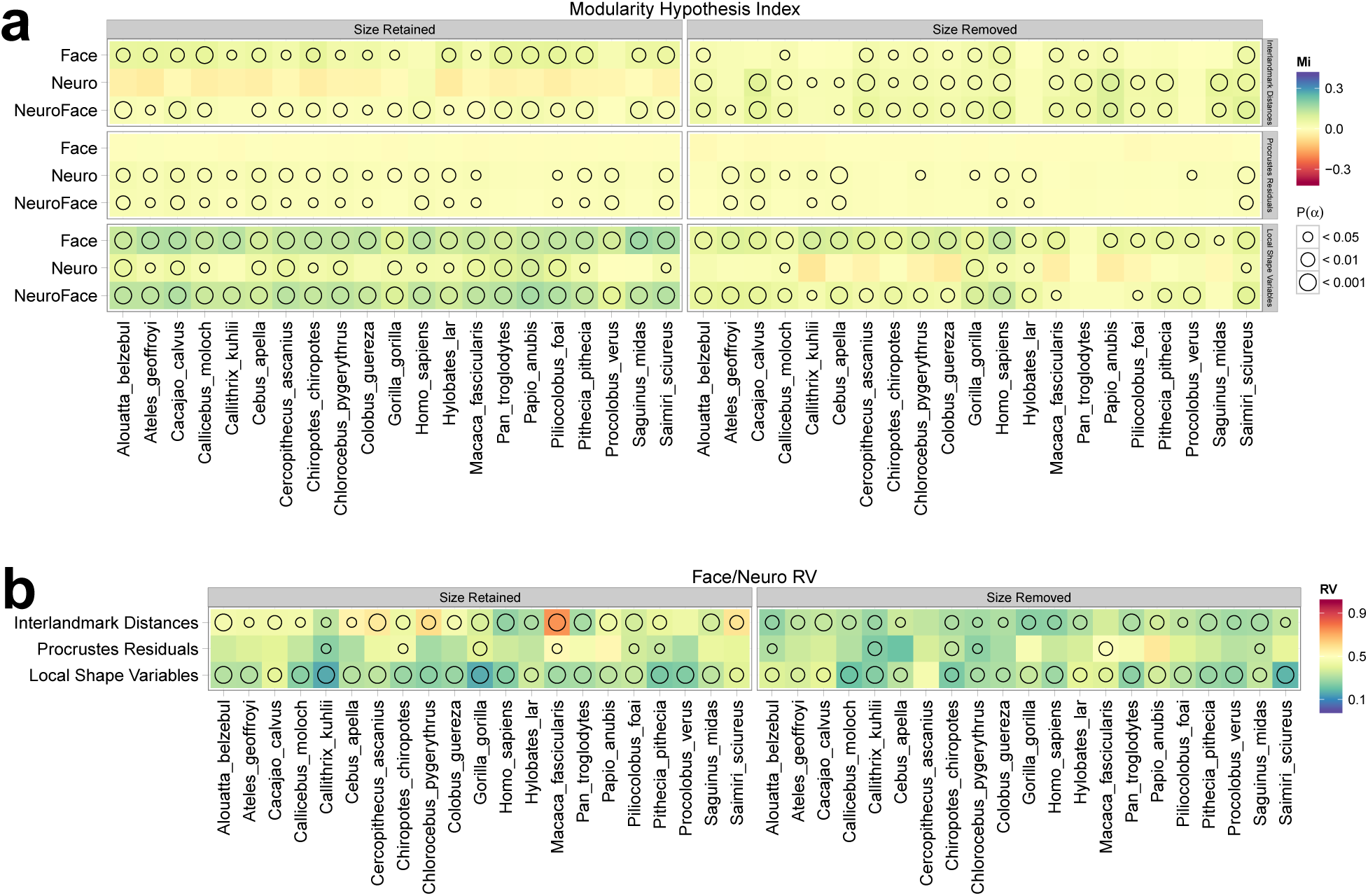
MHI (a) and RV (b) values for regional trait sets. Circles indicate whether a trait set is recognized in a given OTU, with *P*(*α*) indicated by the legend. Notice that blue values are associated with higher MHI values and lower RV values, as the alternate hypothesis for each statistic is formulated in a corresponding manner; see text for details.

Testing the distinction between Face and Neurocranium using RV coefficients (Figure 2b) 364 show that in most cases both regions are considered distinct and valid variational modules for interlandmark distances and local shape variables; for Procrustes residuals, only a handful of taxa show the same result. In this case the correspondence with localized trait sets (Figure 1b) is more difficult due to the lack of independent tests for each region.

### Error Rates

Comparing the distributions of MHI and RV values from theoretical matrices with respect to their structure (Figure 3) shows marked differences between metrics and morphometric variables from which correlations are sampled. In general, the distribution of MHI values obtained from **C***_r_* matrices is the same, while the distributions for **C***_s_* matrices for this metric are more disjoint from the former distribution in local shape variables than in Procrustes residuals, with interlandmark distances representing an intermediate case, regardless of whether size was retained or removed. For RV values, all distributions overlap to some degree; for local shape variables and interlandmark distances with size retained, either distributions (**C***_r_* and **C***_s_*) overlap to a lesser extent.

**Figure 3:**
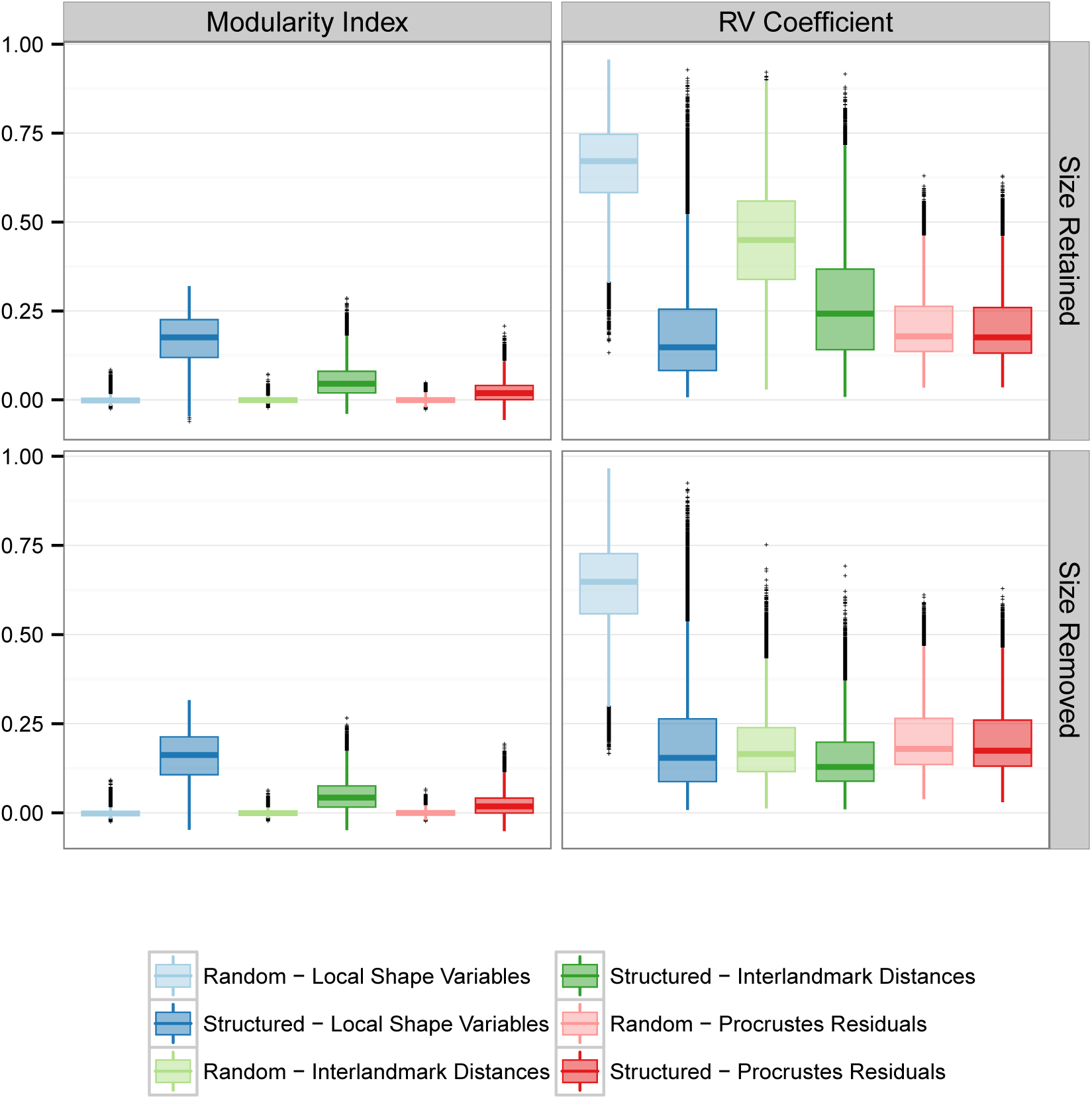
Distribution of Modularity Hypothesis Index and RV Coefficient for theoretical correlation matrices.

Regarding the relationship between significance levels and Type I error rates estimated over **C***_r_* matrices, Figure 4 shows that these quantities approach an identity relationship very closely regardless of whether we use MHI or RV to quantify variational modularity; even at low sample sizes Type I error rates are very close to significance levels. Furthermore, the effect of sampling correlations from size-free distributions does not change Type I error rates.

**Figure 4:**
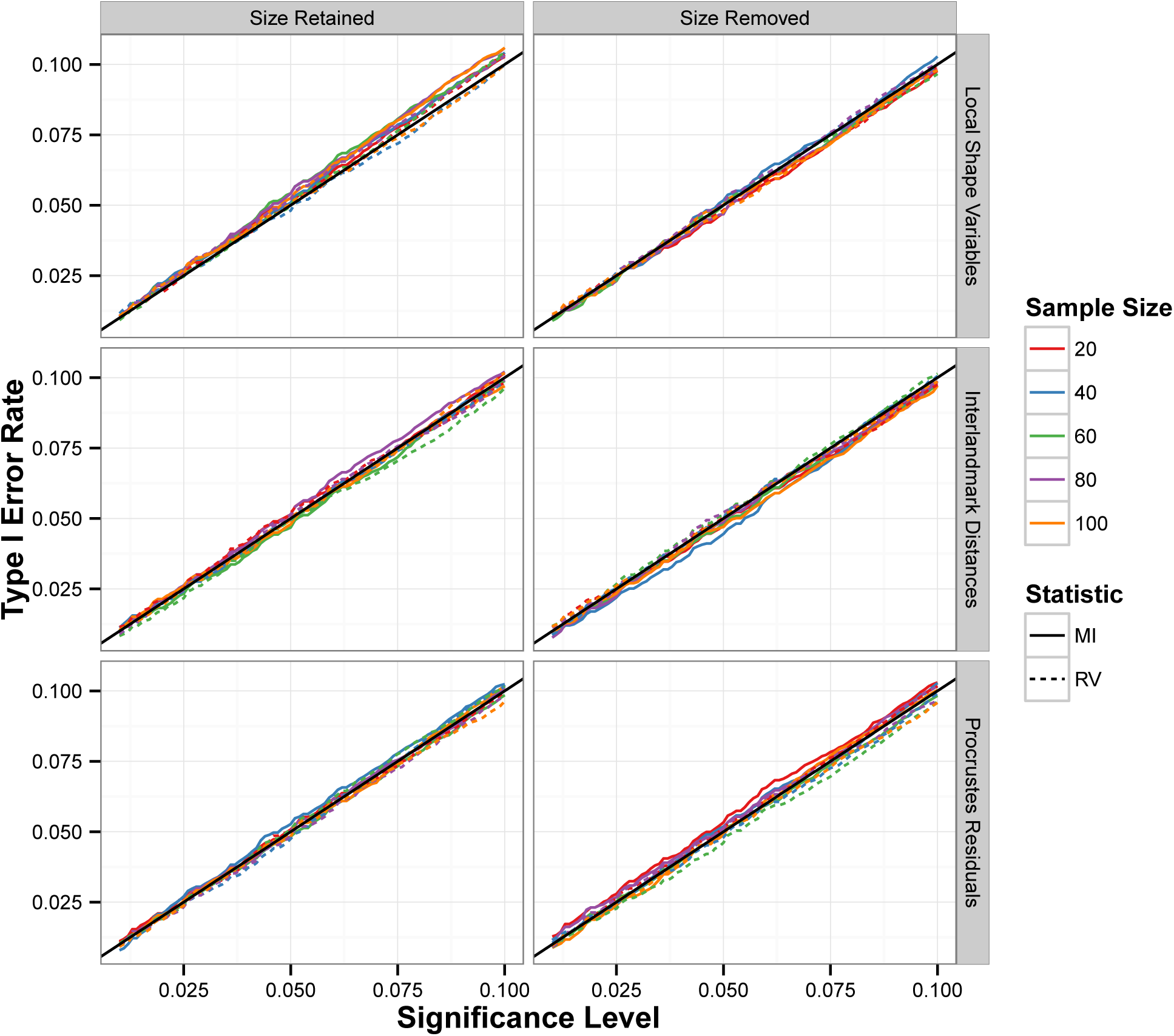
Type I error rates as a function of the chosen significance level regarding tests for variational modularity applied on **C***_r_* correlation matrices. The solid black line represents the identity relationship.

The relationship between power and significance levels (estimated over **C***_s_* matrices) shows substantial differences with respect to the chosen metric (MHI or RV) and to the type of variable that provides sampled correlations. Considering local shape variables (Figure 5), tests using either MHI and RV have high power, even at low sample or effect sizes; increasing these quantities further increases power. However, for lower effect sizes (represented by high average squared correlation between sets, *b*^2^) power for tests using MHI is higher than for those using RV; as effect size increases (lower *b^2^* values), the difference in power between the two statistics decreases. For local shape variables, sampling from its associated size-free correlation distribution implies minor differences in power for both statistics.

**Figure 5:**
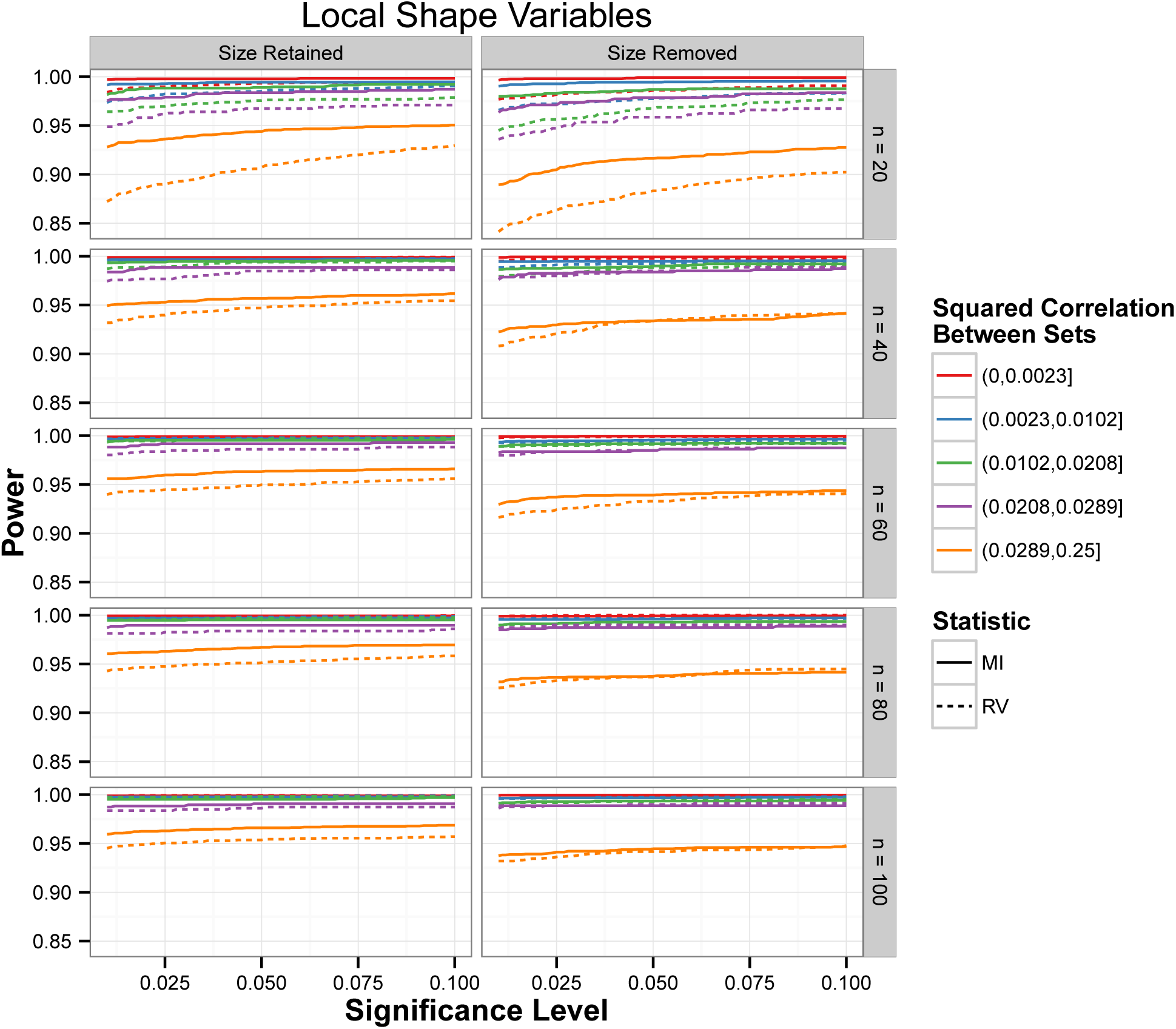
Power for both MHI and RV statistics as a function of the chosen significance levels with respect to tests for variational modularity applied on **C***_s_* matrices with values sampled from the distribution of correlations between local shape variables. Lines are colored with respect to quantiles of the *b^2^* distribution, according to the legend.

For interlandmark distances (Figure 6) there are substantial differences on the relationship between power and significance level if we consider the different parameters. In general, power for tests using MHI are always higher than for tests using RV; this effect is more pronounced on **C***_s_* matrices derived from size-free interlandmark distances, although tests on these matrices have a substantial decrease in power for either tests. However, this decrease is more pronounced for tests based on the RV statistic, since for lower effect sizes (high *b*^2^ values) power approaches an identity relationship with significance level. Sample size also interferes with this relationship since increasing this quantity also increases power when higher effect sizes (low *b*^2^ values) are considered.

**Figure 6:**
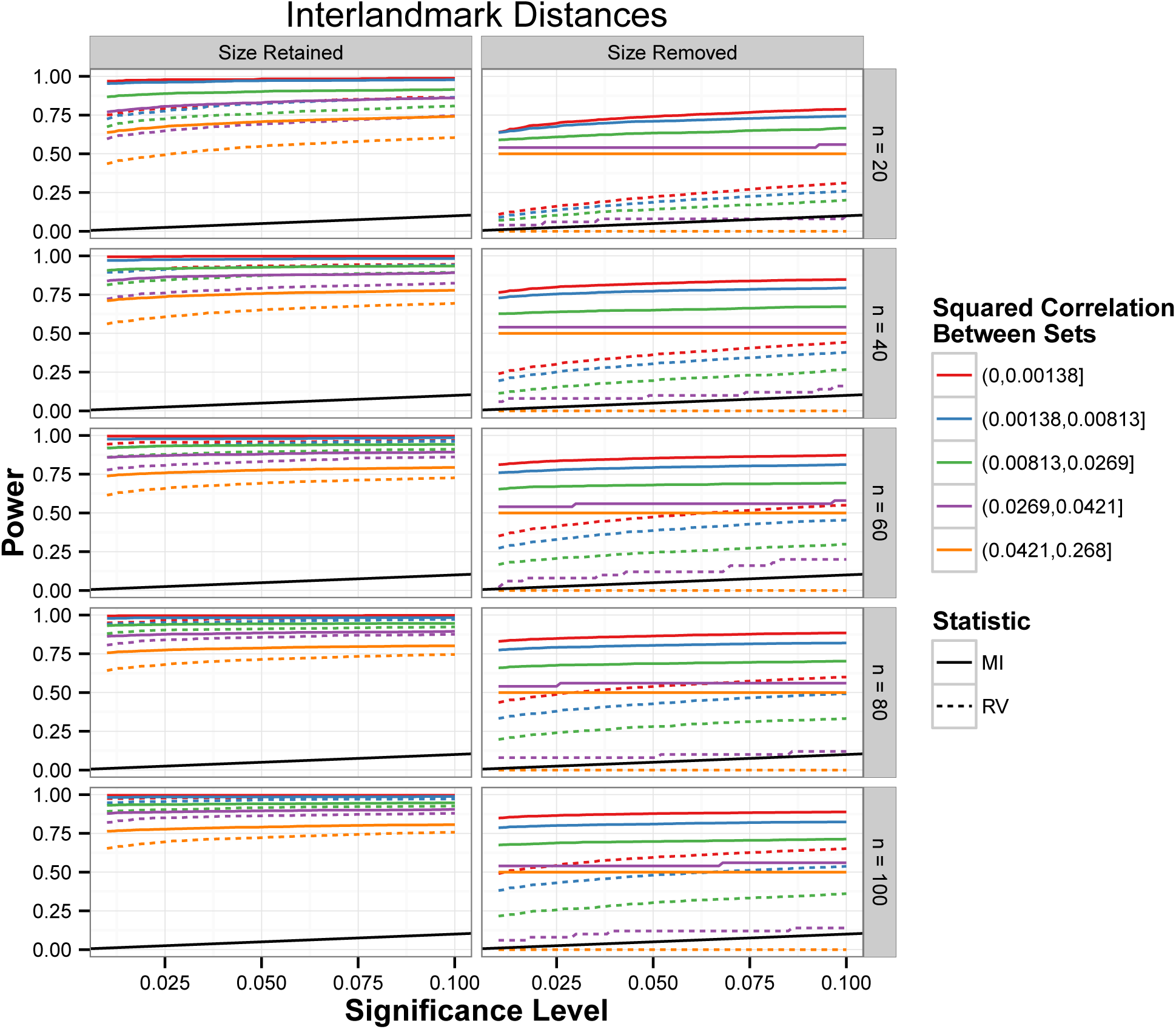
Power for both MHI and RV statistics as a function of the chosen significance levels with respect to tests for variational modularity applied on **C***_s_* correlation matrices with values sampled from the distribution of correlations between interlandmark distances. Lines are colored with respect to quantiles of the *b*^2^ distribution, according to the legend. The solid black line represents the identity relationship.

With respect to Procrustes residuals (Figure 7), tests using either MHI or RV have reduced power regardless of effect or sample size. Sampling from size-free correlation distributions to build **C***_s_* matrices also has little effect. In this case, power for tests performed using RV values approaches an identity relationship with significance level; increasing sample size has some effect, but it does not increase power above 50% in any case.

**Figure 7:**
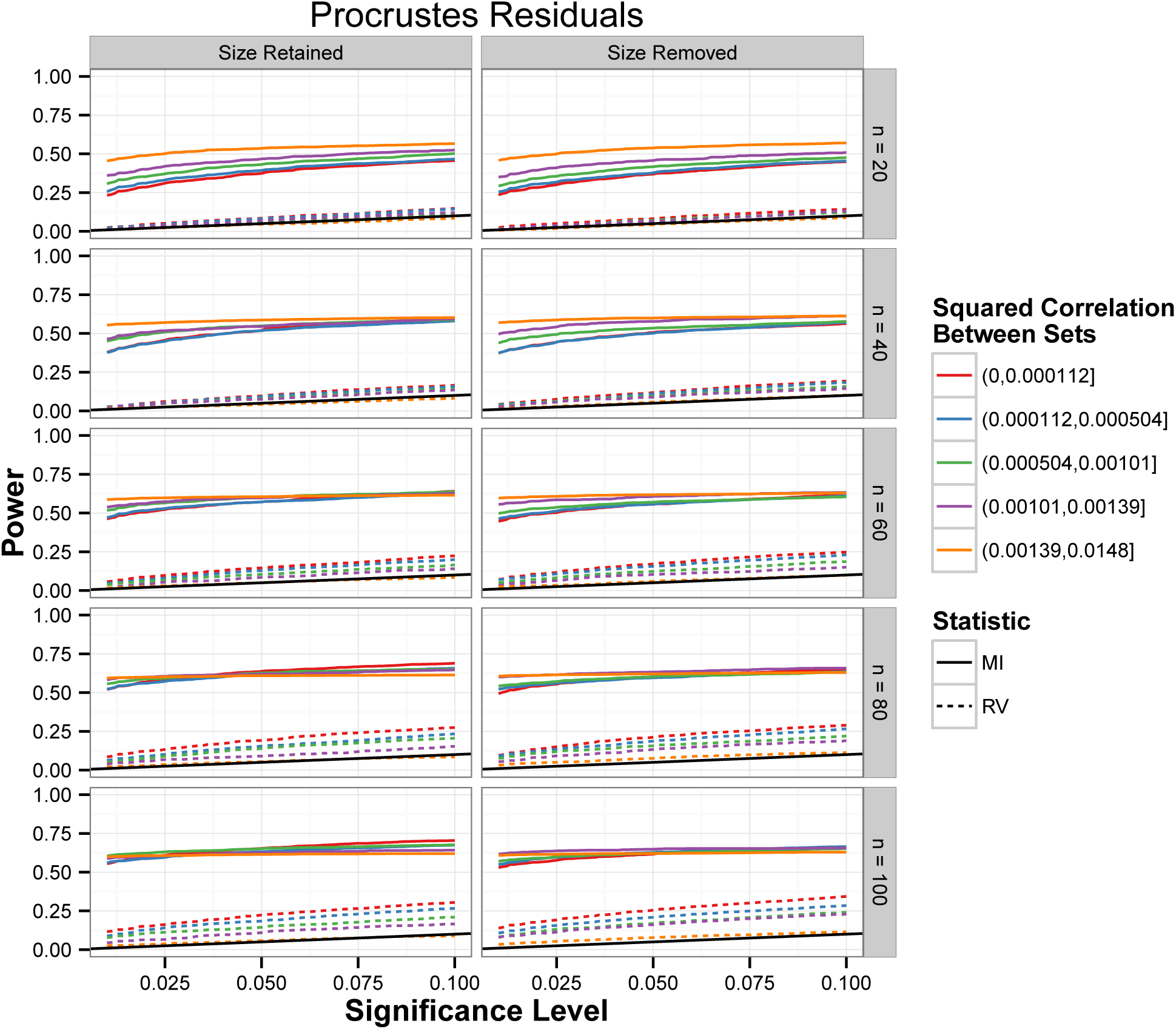
Power for both MHI and RV statistics as a function of the chosen significance levels with respect to tests for variational modularity applied on **C***_s_* matrices with values sampled from the distribution of correlations between Procrustes residuals. Lines are colored with respect to quantiles of the *b*^2^ distribution, according to the legend. The solid black line represents the identity relationship.

## Discussion

Covariance matrices derived from morphological traits are supposed to represent the pattern of codependence that arises due to a variety of interactions among developing morphological elements (Olson & Miller, 1958; Cheverud, 1996). Such interactions are the expression of local developmental factors, as they interact with the expression profiles of surrounding cell types, producing coordinated changes in their cycles and differentiation, thus integrating resulting tissues in the adult population. Although these events of local integration overlap, and the composed effect over adult covariance patterns may be confusing (Hallgrímsson & Lieberman, 2008; Hallgrímsson *et al*., 2009), we believe that a careful comparison of different yet equally proper ways of measuring and representing form may be informative of the underlying processes that produce covariances.

Due to the minimization of quadratic distances among homologous landmarks during GPA, covariance matrices derived from Procrustes estimators lose the signal of localized effects on covariance patterns (Linde & Houle, 2009). Therefore, the use of Procrustes estimators to investigate morphological integration or modularity implicitly implies in a divorce between the phenomenom we would like to investigate and the representation we choose to use. This disconnection between theory and measurement may have grave consequences for the hypotheses we wish to test (Houle *et al*., 2011); such consequences are observable in both our empirical tests and those tests performed on theoretical matrices, as we explore below.

The mammalian Basicranium originates from thirteen precursor tissues derived from both paraxial mesoderm and neural crest, and they may merge to form single bones, such as the sphenoid (Jiang *et al*., 2002; Lieberman, 2011). Furthermore, these precursors display a mosaic of endochondral and intramembranous ossification early in development, and, as the brain grows afterwards, it induces a pattern of internal resorption and exterior deposition on the underlying posterior Basicranium (Lieberman *et al*., 2000); meanwhile, the anterior portion suffers influence from the development of Facial elements (Bastir & Rosas, 2005). Thus, since the Basicranium ossifies early in development, the composed effect of all posterior steps of cranial development will overshadow any pattern of integration this region might have, as predicted by the palimpsest model of development (Hallgrímsson *et al*., 2009). Moreover, the angulation between anterior and posterior elements of the Basicranium has changed sensibly during primate evolution, and such property appears to have evolved in coordination with Facial growth relative to the cranial Vault, accomodating both structures on each other (Scott, 1958; Lieberman *et al*., 2000, 2008).

Due to this heterogeneity of developmental processes acting on the Basicranium, we would not expect it to be a variational module, further expecting that a test of this property over skull covariance patterns will fail to reject the null hypothesis of random association. However, considering the 42 tests performed over covariance matrices derived from Procrustes residuals regarding localized hypotheses (midpanels of Figure 1a), the Basicranium is detected as a valid variational module in 27 cases, distributed through matrices with size either retained or removed; in some cases (e.g. *Alouatta, Cercopithecus)*, only the Basicranium is detected. Thus, Procrustes residuals show a pattern of detection of variational modules opposite to the expectation for this particular test. In covariances matrices derived from interlandmark distances (upper panels of Figure 1a), the Basicranium is not detected a single time; with local shape variables (lower panels of Figure 1a), the Basicranium is detected only three out of 42 times.

These two remaining types of representations, interlandmark distances and local shape variables, show patterns of detection of variational modules that are both consistent to the expectations derived from developmental and functional interactions and to the patterns emerging from these representations when size variation is removed. Considering that interlandmark distances are on a ratio scale (Houle *et al*., 2011), isometric variation will be represented to a greater extent when compared to subtle allometric relationships, due to the multiplicative nature of biological growth (Huxley, 1932). Therefore, the Oral trait set is detected as a valid variational module, considering covariance matrices among interlandmark distances (upper left panel of Figure 1a), since this region is strongly affected by the induction of bone growth due to muscular activity beginning in the pre-weaning period (Zelditch & Carmichael, 1989; Hallgrímsson *et al*., 2009). Furthermore, in these matrices, allometric interactions are associated with the strength of association between traits in Oral, Nasal, Zygomatic and Vault regions in an evolutionary scale (Chapter 3); thus, allometric relationships also play some role in the determination of covariance/correlation patterns for other subregions, although such pattern may be masked in interlandmark distances by the effect of isometric variation. On the other hand, the patterns expressed in covariance/correlation matrices for local shape variables are influenced by allometric relationships defined on a proper scale (Jolicoeur, 1963; Houle *et al*., 2011; Márquez *et al*., 2012) and local developmental processes. Thus, they reflect this association between integration and allometry.

By removing allometric effects from local shape variables (lower right panel of Figure 1a), variational modularity can still be detected in both Oral and Nasal regions, while in a number of species, the Vault region is no longer detected as a variational module. Vault integration may be achieved through both allometric relationships and the effect of relative brain growth, since Vault elements arise mostly through intramembranous ossification, induced by the secretion of signaling factors from the outer brain tissues, with a modest but necessary contribution of mesoderm-derived tissue that undergoes endochondral ossification (Jiang *et al*., 2002; Rice *et al*., 2003; Franz-Odendaal, 2011; Lieberman, 2011). In humans, the growth of Vault osteological elements occur, from a topological standpoint, without deviations from an isometric growth model, forming regular connections among bones (Esteve-Altava & Rasskin-Gutman, 2014), since the boundary of interactions between the tissue inducing growth (brain) and the tissue whose growth is being induced (Vault bones) is also regular. Thus, as this effect dominates the latter stages of pre-natal development (Hallgrímsson & Lieberman, 2008; Lieberman, 2011), the overall effect of brain growth over skull growth patterns in this stage may mirror an affine transformation, rendering it undetectable over covariance patterns derived from local shape variables, since their estimation excludes shape variation associated with affine transformations (Márquez *et al*., 2012). In contrast, Oral and Nasal elements have more complex patterns of connectivity arising from their tight integration with the soft tissues that compose the remaining elements of the Face (Lieberman, 2011; Esteve-Altava *et al*., 2013; Esteve-Altava & Rasskin-Gutman, 2014), thus producing an intrincate pattern of associations that may be responsible for the variational modularity we are able to detect in all species, regardless of whether size variation is retained or removed.

The differences between the pattern of detection in all representations for local trait sets for RV values (Figure 1b) makes a similar interpretation of the results for this metric difficult, as opposed to the results regarding MHI values. The variance of RV values among species seems to indicate that RV values are sensitive to the magnitude of morphological integration, as *Papio* and *Macaca* are among the Catarrhines who display large values of this quantity (Oliveira *et al*., 2009) and score higher (orange/red) RV values for all local trait sets, at least when we consider interlandmark distances. Such sensitivity might be one of the factors responsible for the low power estimated for the RV metric using theoretical covariance matrices; interestingly enough, this effect of magnitude of integration (which is thought to emerge as a consequence of size variation; Wagner, 1984; Marroig & Cheverud, 2001, 2005; Porto *et al*., 2013) seems to also affect Procrustes residuals regardless of whether covariance structure arising from allometric relationships is removed or not.

However, for the set of hypotheses concerning differences between Facial and Neurocranial traits (Figure 2), there is substantial agreement between tests performed using MHI and RV values. For MHI values (Figure 2a), the overall pattern of detection is similar to the pattern detected in local trait sets, for all variable types; for RV values (Figure 2b), there is ample support for the hypotheses that both Face and Neurocranium represent variational modules, in both local shape variables and interlandmark distances. These regions have marked differences in timing and pattern formation during development, as observed from the behavior of their composing units (Zelditch & Carmichael, 1989; Hallgrímsson *et al*., 2009; Lieberman, 2011; Esteve-Altava & Rasskin-Gutman, 2014); therefore, the more general pattern of distinction between Face and Neurocranium is detected regardless of the metric chosen to represent modularity.

### Theoretical Matrices and Error Rates

The distribution of MHI and RV values obtained from the theoretical matrices (Figure 3) is a starting point for understanding the differences in power for tests using these two metrics (Figures 5–7). For MHI values, the distribution obtained from random (**C***_r_*) matrices is consistently the same, regardless of what representation we used to sample correlations, or whether size was retained or removed. On the other hand, the distribution of RV values for random matrices change depending on the representation sampled or whether size variation has been removed or retained. Moreover, this change in behavior for the distribution of RV values for **C***_r_* matrices implies different levels of overlap between these null distributions and the distribution of values obtained for structured (**C***_s_*) matrices.

For Procrustes residuals, a substantial overlap occurs regardless of whether size variation was removed or retained; unsurprisingly, our estimates of power for RV in this type of representation are very low (Figure 7); nonetheless, power estimated for tests based on MHI is lower than in other types of representation, since the difference between within-set and between-set correlations in Procrustes residuals (Figure S2) is the lowest of all representations. A substantial overlap in RV distributions for **C***_r_* and **C***_s_* matrices also occurs with interlandmark distances when size variation is removed, and it implies in low power for tests using RV values in this type of representation (right column of Figure 6). However, in this case there is a substantial difference between within-set and between-set correlations (Figure S2), and tests using MHI to represent modularity are still able to detect such difference (albeit with reduced power) when compared to tests over **C***_s_* matrices derived from interlandmark distances with size retained (Figure 6). In those cases where both distributions for **C***_r_* and **C***_s_* matrices do not overlap substantially — for example, when local shape variables are considered (Figure 5) — power for tests performed using MHI values is always higher than for tests using RV except when sample sizes are very high; in this case power for both metrics are similar. The same behavior can also be observed in interlandmark distances when size is retained (left column of Figure 6).

These results indicate that RV coefficents are more sensitive to the absolute value of both within-set and between-set correlation distributions than MHI values. For interlandmark distances (Figure 6), removing size variation reduced the average value of both correlation distributions by a similar amount (Figure S2); the difference between average correlations in these two sets actually increases, going from 0.042 to 0.061 when size is removed. However, since the actual average correlations for these two distributions approach zero, tests based on RV lose power more rapidly than tests based on MHI. This sensitivity might be associated to the use of squared covariances, as shown by Equation 4, while Modularity Hypothesis Indexes use correlations directly (Equation 3). Furthermore, as observed by Fruciano et al. (2013), sample sizes sensibly influence the estimation of RV coefficients, and we demostrate here that such sensitivity also extends to estimates of power for tests using this metric.

On the other hand, our estimates of power for tests using MHI indicate that it is more robust to differences in absolute correlation values or sample sizes, thus allowing comparisons across more heterogeneous settings, such as our comparison between different representations of form, with substantial variation in sample sizes. The detection of variational modularity is akin to Student’s *t* test, since we are trying to determine whether two groups of observations (correlations between traits in the same subset *versus* correlations between traits in different subsets) have a significant difference in average values; we use resampling procedures to estimate significance in this case only due to the interdependency between pairwise correlations. Thus, as Modularity Hypothesis Indexes are estimated with a formula that is very similar to that of a *t* statistic (difference between location parameters for two groups, divided by a scale parameter — ICV; Equation 3), we believe that this statistic is a proper way to represent variational modularity; the robustness of the tests using this statistic reinforce this belief.

Our approach for constructing theoretical matrices attempts to simulate the most simple situation, that is, the situation where there are only two subsets of traits, akin to the distinction between Face and Neurocranium in our empirical dataset (Figure 2). In this setting, both statistics are capable of detecting this distinction, except when both are used on covariance/correlation patterns derived from Procrustes residuals. However, even though we built theoretical matrices using correlations sampled from these estimators, actually simulating the interference in covariance structure that such estimators produce in our theoretical matrices is quite difficult. Furthermore, constructing such matrices with more complicated patterns (with three modules, for instance) while maintaining their connection to the correlation distributions of each morphometric type is also difficult, due to the restriction on positive-definiteness we enforce on them. Thus, the lack of differences in type I error rates for all cases may be a limitation of our scheme for building theoretical matrices.

The issues we found with the use of Procrustes estimators for covariance matrices and the use of RV coefficients to estimate and detect variational modularity may explain results found by other authors; for instance, Martínez-Abadías et al. (2011) has found no evidence that genetic and phenotypic covariance structure for human skulls conforms to a modular structure, since all tests performed by these authors failed to reject the null hypothesis of random association. These authors use Procrustes estimators to represent covariance structure, and test their hypothesis of partitioning (Face, Vault and Base) using RV as the statistic representing variational modularity. Since this combination implies in very low power (Figure 7), not rejecting the null hypothesis in their case might be a consequence of the choice of estimates of both covariance structure and variational modularity; thus, these authors’ assertion of pervasive genetic integration in the human skull may be misleading, considering that skull covariance patterns in humans are one of the most modular examples of such patterns when compared to other mammals (Porto *et al*., 2009) or catarrhine primates (Oliveira *et al*., 2009).

The approach we explore in the present work is but one of the different ways one can investigate the association between genetic, functional and developmental interactions and correlation structure (Mitteroecker & Bookstein, 2007). For example, Perez et al. (2009) relies on abstracting correlation matrices into networks, then using community-detection algorithms to search for modular patterns without *a priori* hypotheses, associating their results with knockout experiments that support the communities they found among traits; however, it is not clear how much relevant information is retained in these network representations. Furthermore, the authors use Procrustes estimators, which may bias the detection of modularity patterns in this setting in the same manner as we demonstrated here.

Monteiro et al. (2005) assumes that the underlying morphogenetic components of the rodent mandible behave as modules, further investigating the patters of correlation between these units in both within-species variation and between-species variation among Echimids; Monteiro & Nogueira (2010), relying on the correspondence of these units through mammalian diversification did the same to phylostomid bats. Although using a landmark-based approach to represent morphological variation, the authors do not use Procrustes estimators to represent covariance structure among these units, and the pattern of reorganization of correlation structure among these units associated with niche diversification in phylostomids seems robust, considering that this radiation may have been associated with a very heterogeneous adaptive landscape, and such heterogeneity may lead to a reorganization of correlation patterns (Jones, 2007; Jones *et al*., 2012; Melo & Marroig, 2015).

Another valid approach is to model certain aspects of development as null hypotheses; Esteve-Altava & Rasskin-Gutman (2014), investigating the pattern of connections among human cranial bones, conceived the null hypothesis that unconstrained bone growth is sufficient to explain the observed patterns. Such approach could be extended to investigate morphological covariance structure; if we consider the geometric properties of the features we measure (Mitteroecker & Bookstein, 2007), one could formulate the null hypothesis that topological proximity is a sufficient explanation for the observed covariance structure, against the alternative hypothesis that local developmental processes coupled with functional interactions produce stronger relationships among close elements that surpass these purely topological interactions. Alternatively, one could actively look for variational boundaries between regions, as boundary formation is a phenomenom that has been well studied under a dynamical perspective on development (e.g. Turing, 1952; Meinhardt, 1983; Tiedemann *et al*., 2012).

The approach of partitioning covariance matrices into blocks that correspond to inferred modular associations has the advantage that it is simple from an operational standpoint; however, modularity patterns are almost certainly not expressed in phenotypic data as the binary hypotheses we used here (Hallgrímsson *et al*., 2009). Thus, hypotheses and inferences made from them have to be contextualized in the light of developmental dynamics, since the measurements we make and the parameters we estimate have to be properly connected to the models we are considering; otherwise, inferences made from such models may be devoid of meaning (Wagner, 2010; Houle *et al*., 2011).

## Conclusion

Here we show that Procrustes estimators for covariance matrices fail to capture the modularity patterns embedded in phenotypic data, regardless of which metric is chosen to represent such patterns, although the combination of this type of variable with RV coefficients for investigating modularity has even more problems than either has alone. Both interlandmark distances and local shape variables seem valid options to represent morphological variation, if their limitations are taken into consideration. We wish to stress this point: any representation of morphological variation has limitations since they are themselves models — at the very least of what it is important to represent — not fully capturing the phenomena we may be interested in.

## Acknowledgements

We thank G. Burin, D. Melo, and A. Porto for comments on an early draft. This work has been funded by CNPq (Conselho Nacional de Pesquisa e Desenvolvimento Tecnológico) and FAPESP (Fundação de Apoio à Pesquisa do Estado de São Paulo).

## References

Adams, D.C. & Otárola-Castillo, E. 2013. geomorph: an r package for the collection and analysis of geometric morphometric shape data. Methods in Ecology and Evolution 4: 393–399.

Adams, D.C., Rohlf, F.J. & Slice, D.E. 2004. Geometric morphometrics: Ten years of progress following the “revolution”. Italian Journal of Zoology 71: 5–16.

Andrade, R.F.S., Rocha-Neto, I.C., Santos, L.B.L., Santana, C.N. de, Diniz, M.V.C. & Lobão, T.P. et al. 2011. Detecting Network Communities: An Application to Phylogenetic Analysis. PLoS Computational Biology 7: e1001131.

Arnold, S.J., Bürger, R., Hohenlohe, P.A., Ajie, B.C. & Jones, A.G. 2008. Understanding the evolution and stability of the G-matrix. Evolution 62: 2451–2461.

Bastir, M. & Rosas, A. 2005. Hierarchical nature of morphological integration and modularity in the human posterior face. American Journal of Physical Anthropology 128: 26–34.

Bookstein, F.L. 1982. Foundations of Morphometrics. Annual Review of Ecology and Systematics 13: 451–470.

Bookstein, F.L. 1991. Morphometric tools for landmark data: geometry and biology. Cambridge University Press, Cambridge.

Bookstein, F.L. 1989. Principal warps: Thin plate splines and the decomposition of deformations. IEEE Transactions on Pattern Analysis and Machine Intelligence 11: 567–585.

Bookstein, F.L., Chernoff, B., Elder, R., Humphries, Smith, G. & Strauss, R. 1985. Morphometrics in Evolutionary Biology. The Academy of Natural Sciences of Philadelphia, Philadelphia.

Cardini, A. & Polly, P.D. 2013. Larger mammals have longer faces because of size-related constraints on skull form. Nature Communications 4.

Cheverud, J.M. 1996. Developmental integration and the evolution of pleiotropy. American Zoology 36: 44–50.

Cheverud, J.M. 1982. Phenotypic, genetic, and environmental morphological integration in the cranium. Evolution 36: 499–516.

Cheverud, J.M. & Richtsmeier, J.T. 1986. Finite-Element Scaling Applied to Sexual Dimorphism in Rhesus Macaque (Macaca Mulatta) Facial Growth. Systematic Biology 35: 381–399.

Cheverud, J.M., Kohn, L.A.P., Konigsberg, L.W. & Leigh, S.R. 1992. Effects of fronto-occipital artificial cranial vault modification on the cranial base and face. American Journal of Physical Anthropology 88: 323–345.

Cheverud, J.M., Routman, E.J. & Irschick, D.J. 1997. Pleiotropic Effects of Individual Gene Loci on Mandibular Morphology. Evolution 51: 2006–2016.

Cheverud, J.M., Wagner, G.P. & Dow, M.M. 1989. Methods for the comparative analysis of variation patterns. Evolution 38: 201–213.

Drake, A.G. & Klingenberg, C.P. 2010. Large Scale Diversification of Skull Shape in Domestic Dogs: Disparity and Modularity. The American Naturalist 175: 289–301.

Escoufier, Y. 1973. Le Traitement des Variables Vectorielles. Biometrics 29: 751–760.

Espinosa-Soto, C. & Wagner, A. 2010. Specialization can drive the evolution of modularity. PLoS Comput. Biol. 6: e1000719.

Esteve-Altava, B. & Rasskin-Gutman, D. 2014. Beyond the functional matrix hypothesis: a network null model of human skull growth for the formation of bone articulations. Journal of Anatomy 225: 306–316.

Esteve-Altava, B., Marugán-Lobón, J., Botella, H., Bastir, M. & Rasskin-Gutman, D. 2013. Grist for Riedl’s mill: A network model perspective on the integration and modularity of the human skull. Journal of Experimental Zoology Part B: Molecular and Developmental Evolution 320: 489–500.

Falconer, D.S. & Mackay, T.F.C. 1996. Introduction to Quantitative Genetics, 4th ed. Addison Wesley Longman, Harlow, Essex.

Fortuna, M.A., García, C., Guimarães Jr., P.R. & Bascompte, J. 2008. Spatial mating networks in insect-pollinated plants. Ecology Letters 11: 490–498.

Franz-Odendaal, T.A. 2011. Epigenetics in Bone and Cartilage Development. In: Epigenetics: Linking Genotype and Phenotype in Development andEvolution (B. Hallgrímsson & B. K. Hall, eds), pp. 195–220. University of California Press.

Fruciano, C., Franchini, P. & Meyer, A. 2013. Resampling-Based Approaches to Study Variation in Morphological Modularity. PLoS ONE 8: e69376.

Genini, J., Morellato, L.P.C., Guimarães Jr., P.R. & Olesen, J.M. 2010. Cheaters in mutualism networks. Biology Letters 6: 494–497.

Goodall, C. 1991. Procrustes methods in the statistical analysis of shape. Journal of the Royal Statistical Society. Series B (Methodological) 53: 285–339.

Goswami, A. & Polly, P.D. 2010. The influence of modularity on cranial morphological disparity in Carnivora and Primates (Mammalia). PLoS ONE 5: e9517.

Grabowski, M.W., Polk, J.D. & Roseman, C.C. 2011. Divergent patterns of integration and reduced constraint in the human hip and the origins of bipedalism. Evolution 65: 1336–1356.

Haber, A. 2015. The Evolution of Morphological Integration in the Ruminant Skull. Evolutionary Biology 42: 99–114.

Hallgrímsson, B. & Lieberman, D.E. 2008. Mouse models and the evolutionary developmental biology of the skull. Integrative and Comparative Biology 48: 373–384.

Hallgrímsson, B., Jamniczky, H., Young, N.M., Rolian, C., Parsons, T.E. & Boughner, J.C. et al. 2009. Deciphering the Palimpsest: Studying the Relationship Between Morphological Integration and Phenotypic Covariation. Evolutionary Biology 36: 355–376.

Herring, S.W. 2011. Muscle-Bone Interactions and the Development of Skeletal Phenotype. In: Epigenetics: Linking Genotype and Phenotype in Development andEvolution (B. Hallgrímsson & B. K. Hall, eds), pp. 221–237. University of California Press.

Houle, D., Pélabon, C., Wagner, G.P. & Hansen, T.F. 2011. Measurement and Meaning In Biology. The Quartely Review of Biology 86: 3–34.

Huckemann, S. 2011. Inference on 3D Procrustes Means: Tree Bole Growth, Rank Deficient Diffusion Tensors and Perturbation Models: Inference on 3D Procrustes means. Scandinavian Journal of Statistics no–no.

Huckemann, S.F. 2012. On the meaning of mean shape: manifold stability, locus and the two sample test. Annals of the Institute of Statistical Mathematics 64: 1227–1259.

Huxley, J.S. 1932. Problems of relative growth.

Jiang, X., Iseki, S., Maxson, R.E., Sucov, H.M. & Morriss-Kay, G.M. 2002. Tissue Origins and Interactions in the Mammalian Skull Vault. Developmental Biology 241: 106–116.

Jolicoeur, P. 1963. The Multivariate Generalization of the Allometry Equation. Biometrics.

Jones, A.G. 2007. The mutation matrix and the evolution of evolvability. Evolution 61: 727–745.

Jones, A.G., Bürger, R., Arnold, S.J., Hohenlohe, P.A. & Uyeda, J.C. 2012. The effects of stochastic and episodic movement of the optimum on the evolution of the G-matrix and the response of the trait mean to selection. Journal of evolutionary biology 1–22.

Kendall, D.G. 1984. Shape manifolds, procrustean metrics, and complex projective spaces. Bulletin of the London Mathematical Society 16: 81–121.

Kent, J.T. & Mardia, K.V. 1997. Consistency of Procrustes Estimators. Journal of the Royal Statistical Society: Series B (Statistical Methodology) 59: 281–290.

Klingenberg, C.P. 2011. MorphoJ: an integrated software package for geometric morpho-metrics. Molecular Ecology Resources 11: 353–357.

Klingenberg, C.P. 2009. Morphometric integration and modularity in configurations of landmarks: tools for evaluating a priori hypotheses. Evolution & Development 11: 405–421.

Klingenberg, C.P. & Leamy, L.J. 2001. Quantitative genetics of geometric shape in the mouse mandible. Evolution 55: 2342–2352.

Klingenberg, C.P., Barluenga, M. & Meyer, A. 2002. Shape analysis of symmetric structures: Quantifying variation among individuals and asymmetry. Evolution 56: 1909–1920.

Klingenberg, C.P., Leamy, L.J. & Cheverud, J.M. 2004. Integration and Modularity of Quantitative Trait Locus Effects on Geometric Shape in the Mouse Mandible. Genetics 166: 1909–1921.

Lele, S. 1993. Euclidean distance matrix analysis (EDMA): estimation of mean form and mean form difference. Mathematical Geology 25: 573–602.

Lele, S.R. & McCulloch, C.E. 2002. Invariance, Identifiability, and Morphometrics. Journal of the American Statistical Association 97: 796–806.

Lieberman, D.E. 2011. Epigenetic Integration, Complexity and Evolvability of the Head: Rethinking the Functional Matrix Hypothesis. In: Epigenetics: Linking Genotype and Phenotype in Development and Evolution (B. Hallgrímsson & B. K. Hall, eds), pp. 271–289. University of California Press.

Lieberman, D.E., Hallgrímsson, B., Liu, W., Parsons, T.E. & Jamniczky, H.A. 2008. Spatial packing, cranial base angulation, and craniofacial shape variation in the mammalian skull: testing a new model using mice. Journal of Anatomy 212: 720–735.

Lieberman, D.E., Ross, C.E & Ravosa, M.J. 2000. The primate cranial base: Ontogeny, function, and integration. American Journal of Physical Anthropology 113: 117–169.

Linde, K. van der & Houle, D. 2009. Inferring the Nature of Allometry from Geometric Data. Evolutionary Biology 36: 311–322.

Lynch, M. & Walsh, B. 1998. Genetics and analysis of quantitative traits. Sinauer Associates, Sunderland.

Mantel, N. 1967. The detection of disease clustering and a generalized regression approach. Cancer Res 27: 209–220.

Marcucio, R.S., Cordero, D.R., Hu, D. & Helms, J.A. 2005. Molecular interactions coordinating the development of the forebrain and face. Developmental Biology 284: 48–61.

Marroig, G. & Cheverud, J.M. 2001. A comparison of phenotypic variation and covariation patterns and the role of phylogeny, ecology, and ontogeny during cranial evolution of new world monkeys. Evolution 55: 2576–2600.

Marroig, G. & Cheverud, J.M. 2005. Size as a line of least evolutionary resistance: Diet and adaptive morphological radiation in new world monkeys. Evolution 59: 1128–1142.

Marroig, G. & Cheverud, J.M. 2010. Size as a line of least resistance II: direct selection on size or correlated response due to constraints? Evolution 64: 1470–1488.

Martínez-Abadías, N., Esparza, M., vold, T. Sjø, González-José, R., Hernández, M. & Klingenberg, C.P. 2011. Pervasive genetic integration directs the evolution of human skull shape. Evolution 66: 1010–1023.

Márquez, E.J., Cabeen, R., Woods, R.P. & Houle, D. 2012. The Measurement of Local Variation in Shape. Evolutionary Biology 39: 419–439.

Meinhardt, H. 1983. A boundary model for pattern formation in vertebrate limbs. Journal of Embryology and Experimental Morphology 76: 115–137.

Meinhardt, H. 2008. Models of biological pattern formation: from elementary steps to the organization of embryonic axes. Current topics in developmental biology 81: 1–63.

Melo, D. & Marroig, G. 2015. Directional selection can drive the evolution of modularity in complex traits. Proceedings of the National Academy of Sciences 112: 470–475.

Minelli, A. 2011. A principle of developmental inertia. Epigenetics: Linking Genotype and Phenotype in Development and Evolution 116–133.

Mitteroecker, P. & Bookstein, F.L. 2007. The conceptual and statistical relationship between modularity and morphological integration. Systematic Biology 56: 818–836.

Mitteroecker, P., Gunz, P., Bernhard, M., Bookstein, F.L. & Schaefer, K. 2004. Comparison of cranial ontogenetic trajectories among great apes and humans. Journal of Human Evolution 46: 679–697.

Monteiro, L.R. & Nogueira, M.R. 2010. Adaptive radiations, ecological specialization, and the evolutionary integration of complex morphological structures. Evolution 64: 724–744.

Monteiro, L.R., Bonato, V. & Reis, S.F. 2005. Evolutionary integration and morphological diversification in complex morphological structures: mandible shape divergence in spiny rats (Rodentia, Echimyidae). Evolution & Development 7: 429–439.

Newman, M.E.J. 2006. Modularity and community structure in networks. Proceedings of the National Academy of Sciences 103: 8577–8582.

Neyman, J. & Scott, E.L. 1948. Consistent Estimates Based on Partially Consistent Observations. Econometrica 16: 1–32.

Oliveira, F.B., Porto, A. & Marroig, G. 2009. Covariance structure in the skull of Catarrhini: a case of pattern stasis and magnitude evolution. Journal of Human Evolution 56: 417–430.

Olson, E. & Miller, R. 1958. Morphological integration. University of Chicago Press, Chicago.

Pearson, K. & Davin, A.G. 1924. On the Biometric Constants of the Human Skull. Biometrika 16: 328–363.

Perez, S.I., Aguiar, M.A.M., Guimarães Jr., P.R. & Reis, S.F. dos. 2009. Searching for Modular Structure in Complex Phenotypes: Inferences from Network Theory. Evolutionary Biology, doi: 10.1007/s11692-009-9074-7.

Pélabon, C., Bolstad, G.H., Egset, C.K., Cheverud, J.M., Pavlicev, M. & Rosenqvist, G. 2013. On the Relationship between Ontogenetic and Static Allometry. The American Naturalist 181: 195–212.

Polly, P.D. 2008. Developmental Dynamics and G-Matrices: Can Morphometric Spaces be Used to Model Phenotypic Evolution? Evolutionary Biology 35: 83–96.

Porto, A., Oliveira, F.B., Shirai, L.T., Conto, V. de & Marroig, G. 2009. The evolution of modularity in the mammalian skull I: morphological integration patterns and magnitudes. Evolutionary Biology 36: 118–135.

Porto, A., Shirai, L.T., Oliveira, F.B. de & Marroig, G. 2013. Size Variation, Growth Strategies, and the Evolution of Modularity in the Mammalian Skull. Evolution 67: 3305–3322.

R Core Team. 2015. R: A Language and Environment for Statistical Computing. R Foundation for Statistical Computing, Vienna, Austria.

Ravasz, E., Somera, A.L., Mongru, D.A., Oltvai, Z.N. & Barabási, A.L. 2002. Hierarchical organization of modularity in metabolic networks. Science 297: 1551–1555.

Rice, D.P.C., Rice, R. & Thesleff, I. 2003. Molecular mechanisms in calvarial bone and suture development, and their relation to craniosynostosis. The European Journal of Orthodontics 25: 139–148.

Rohlf, F.J. & Slice, D. 1990. Extensions of the Procrustes Method for the Optimal Superimposition of Landmarks. Systematic Biology 39: 40–59.

Rueffler, C., Hermisson, J. & Wagner, G.P. 2012. Evolution of functional specialization and division of labor. Proceedings of the National Academy of Sciences 109: E326–E335.

Sanger, T.J., Mahler, D.L., Abzhanov, A. & Losos, J.B. 2012. Roles for modularity and constraint in the evolution of cranial diversity among Anolis lizards. Evolution 66: 1525–42.

Schluter, D. 1996. Adaptive radiation along genetic lines of least resistance. Evolution 50: 1766–1774.

Scott, J.H. 1958. The cranial base. American Journal of Physical Anthropology 16: 319–348.

Shirai, L.T. & Marroig, G. 2010. Skull modularity in neotropical marsupials and monkeys: size variation and evolutionary constraint and flexibility. Journal of experimental zoology. Part B, Molecular and developmental evolution 314B: 663–683.

Theobald, D.L. & Wuttke, D.S. 2006. Empirical Bayes hierarchical models for regularizing maximum likelihood estimation in the matrix Gaussian Procrustes problem. Proceedings of the National Academy of Sciences 103: 18521–18527.

Tiedemann, H.B., Schneltzer, E., Zeiser, S., Hoesel, B., Beckers, J. & Przemeck, G.K.H. et al. 2012. From dynamic expression patterns to boundary formation in the presomitic mesoderm. PLoS computational biology 8: e1002586.

Turing, A.M. 1952. The Chemical Basis of Morphogenesis. Philosophical Transactions of the Royal Society of London 237: 37–72.

Wagner, G.P. 1996. Homologues, natural kinds and the evolution of modularity. The American Zoologist 36: 36–43.

Wagner, G.P. 1984. On the eigenvalue distribution of genetic and phenotypic dispersion matrices: evidence for a nonrandom organization of quantitative character variation. Journal of Mathematical Biology 21: 77–95.

Wagner, G.P. 2010. The Measurement Theory of Fitness. Evolution 64: 1358–1376.

Wagner, G.P. & Altenberg, L. 1996. Perspective: complex adaptations and the evolution of evolvability. Evolution 50: 967–976.

Wagner, G.P., Pavlicev, M. & Cheverud, J.M. 2007. The road to modularity. Nature reviews. Genetics 8: 921–931.

Walker, J.A. 2000. Ability of Geometric Morphometric Methods to Estimate a Known Covariance Matrix. Systematic Biology 49: 686–696.

Willmore, K.E., Roseman, C.C., Rogers, J., Cheverud, J.M. & Richtsmeier, J.T. 2009. Comparison of Mandibular Phenotypic and Genetic Integration between Baboon and Mouse. Evolutionary Biology 36: 19–36.

Woods, R.P. 2003. Characterizing volume and surface deformations in an atlas framework: theory, applications, and implementation. NeuroImage 18: 769–788.

Young, N.M. & Hallgrímsson, B. 2005. Serial homology and the evolution of mammalian limb covariation structure. Evolution 59: 2691–2704.

Young, N.M., Wagner, G.P. & Hallgrímsson, B. 2010. Development and the evolvability of human limbs. Proceedings of the National Academy of Sciences 107: 3400–3405.

Zelditch, M.L. & Carmichael, A.C. 1989. Ontogenetic variation in patterns of developmental and functional integration in skulls of Sigmodon fulviventer. Evolution 43: 814–824.

Zelditch, M.L. & Swiderski, D.L. 2011. Epigenetic interactions: the developmental route to functional integration. In: Epigenetics: linking genotype and phenotype in development and evolution, pp. 290–316.

Zelditch, M.L., Swiderski, D.L., Sheets, H.D. & Fink, W.L. 2004. Geometric Morphometrics for Biologists: A Primer, 1st ed. Elsevier.

Zelditch, M.L., Wood, A.R. & Swiderski, D.L. 2009. Building Developmental Integration into Functional Systems: Function-Induced Integration of Mandibular Shape. Evolutionary Biology 36: 71–87.

